# Fast quantitative analysis of timsTOF PASEF data with MSFragger and IonQuant

**DOI:** 10.1101/2020.03.19.999334

**Authors:** Fengchao Yu, Sarah E. Haynes, Guo Ci Teo, Dmitry M. Avtonomov, Daniel A. Polasky, Alexey I. Nesvizhskii

## Abstract

Ion mobility brings an additional dimension of separation to liquid chromatography-mass spectrometry, improving identification of peptides and proteins in complex mixtures. A recently introduced timsTOF mass spectrometer (Bruker) couples trapped ion mobility separation to time-of-flight mass analysis. With the parallel accumulation serial fragmentation (PASEF) method, the timsTOF platform achieves promising results, yet analysis of the data generated on this platform represents a major bottleneck. Currently, MaxQuant and PEAKS are most commonly used to analyze these data. However, due to the high complexity of timsTOF PASEF data, both require substantial time to perform even standard tryptic searches. Advanced searches (e.g. with many variable modifications, semi- or non-enzymatic searches, or open searches for post-translational modification discovery) are practically impossible. We have extended our fast peptide identification tool MSFragger to support timsTOF PASEF data, and developed a label-free quantification tool, IonQuant, for fast and accurate 4-D feature extraction and quantification. Using a HeLa data set published by Meier et al. (2018), we demonstrate that MSFragger identifies significantly (∼30%) more unique peptides than MaxQuant (1.6.10.43), and performs comparably or better than PEAKS X+ (∼10% more peptides). IonQuant outperforms both in terms of number of quantified proteins while maintaining good quantification precision and accuracy. Runtime tests show that MSFragger and IonQuant can fully process a typical two-hour PASEF run in under 70 minutes on a typical desktop (6 CPU cores, 32 GB RAM), significantly faster than other tools. Finally, through semi-enzymatic searching, we significantly increase the number of identified peptides. Within these semi-tryptic identifications, we report evidence of gas-phase fragmentation prior to MS/MS analysis.

## Introduction

A major challenge to identification and quantification of proteins from tissue or cultured cells is the immense complexity of the peptide mixtures that result from enzymatic preparation of these samples for liquid chromatography-mass spectrometry (LC-MS) analysis. Ion mobility spectrometry brings an additional dimension of separation to LC-MS proteomics, significantly improving peptide identification. Following electrospray ionization, ion mobility differentiates gas-phase peptide ions by their size and charge prior to mass analysis. Ion mobility separation occurs on the millisecond timescale, improving selectivity without adding to analysis times. Recently, a commercially available instrument that couples trapped ion mobility spectrometry (TIMS) to time-of-flight (TOF) mass analysis (1) has achieved promising depth of coverage, routinely identifying over 6000 proteins from individual 120-minute LC gradients (2, 3).

Owing to the dual TIMS design of this instrument, where the first region is used for storing ions and the second for ion mobility separation, peptides can be continually selected for sequencing with minimal reduction in duty cycle. This data acquisition method has been termed parallel accumulation-serial fragmentation (PASEF) (2, 3). For typical data-dependent acquisition (DDA) measurements, a survey scan is performed, and the N-highest abundance precursor ions are targeted for tandem mass spectrometry (MS/MS) analysis based on their mass-to-charge ratio (m/z) and mobility. Fast quadrupole switching times allow multiple peptide ions to be targeted for fragmentation during a single ion mobility scan. As a target precursor exits the TIMS region, the quadrupole switches to transmit the corresponding m/z determined by the survey scan. Synchronization of the TIMS device and quadrupole mass filter reduces chimeric spectra and enables removal of singly-charged contaminant ions. Additionally, because of the fast acquisition speed (50-200 ms for a full scan), low-abundance precursors can be repeatedly re-targeted to improve MS/MS spectrum quality (2, 3).

A current major limitation of the PASEF proteomics method is long post-acquisition analysis time due to the high dimensionality of the data and large number of acquired MS/MS scans. MaxQuant (4, 5) and PEAKS (6) are both capable of processing PASEF data but require roughly three hours to perform a standard tryptic search given a raw data file from a two hour gradient. Neither MaxQuant nor PEAKS are practical for nonspecific digest searches or open searches (7, 8), which are helpful in discovering post-translational modifications. We have recently introduced a fragment ion indexing method and its implementation in an ultrafast database search tool MSFragger (8). The speed of MSFragger makes it well suited for the analysis of large and complex data sets such as those from timsTOF PASEF. As conversion from Bruker’s raw liquid chromatography-ion mobility-mass spectrometry (LC-IMS-MS) format (.d) to an open, searchable format (.mzML) represents another significant computational challenge (up to 90 minutes per single two-hour LC-MS gradient raw file), we also extended MSFragger to read the raw format directly. Here we demonstrate that MSFragger can now perform peptide identification from raw timsTOF PASEF data in a fraction of the time required by other tools.

A second challenge is related to quantification of timsTOF PASEF data. Due to the added ion mobility dimension, previously developed quantification tools need to be extended to LC-IMS-MS data. In MaxQuant this is done by slicing a 4-D space (ion mobility, m/z, retention time, and intensity) into multiple 3-D sub-spaces (m/z, retention time, and intensity) and tracing peaks within each sub-space (5). Though MaxQuant only uses every third TOF scan in feature detection, it represents a significant fraction of the overall analysis time. Similarly, PEAKS (6) has extended its functionality to support quantification of timsTOF PASEF data, with analysis times similar to those of MaxQuant. To address this challenge, we introduce IonQuant, a tool that takes Bruker’s raw files and database search results as input to perform fast extracted ion chromatogram (XIC)-based quantification. Using spectral data indexing for XIC tracing in retention and ion mobility dimensions, IonQuant requires ∼10 minutes per file on a desktop computer. IonQuant is integrated seamlessly with MSFragger (8) and the Philosopher validation toolkit (9).

Using timsTOF PASEF HeLa data published by Meier et al. (3) and multi-organism mixture data published by Prianichnikov et al. (5), we show the application of MSFragger and IonQuant to measure the analysis speed and quantitative reproducibility across replicate injections, and compare these results to PEAKS and MaxQuant. We demonstrate how more comprehensive (including semi-enzymatic and open) searches with MSFragger enable deep dives in these data, revealing interesting trends and recovering large numbers of peptides missed in the original analysis. Additionally, our pipeline has spectral library building capabilities and is fully compatible with the Skyline environment for subsequent visualization and targeted exploration of the data. Overall, we showcase a fast, flexible, and accurate computational platform for analyzing timsTOF PASEF proteomics data.

## Experimental Procedures

### Experimental Design and Statistical Rationale

We used data from five experimental conditions (25, 50, 100, 150, and 200 ms TIMS accumulation time) published by Meier et al. (3) in the experiments. Each experimental condition has four technical replicates. Meier et al. (3) concluded that the 100 ms accumulation time gave the best identification results. We used these four replicates with 100 ms accumulation time extensively (performing closed tryptic search, closed semi-enzymatic search, open search, and label free quantification comparisons). We also used data generated from a mixture of three organisms (*H. sapiens, S. cerevisiae*, and *E. coli*) published by Prianichnikov et al. (5). There are two experimental conditions (A and B) that contain the following ratios of each organism with respect to one another: 1:1 (*H. sapiens*), 2:1 (*S. cerevisiae*), and 1:4 (*E. coli*). We used this data to evaluate the quantification accuracy of IonQuant. For identification, we estimated the false-discovery rate (FDR) using the target-decoy approach (10, 11). For quantification, we evaluated the quality with coefficient of variation (CV) and Pearson correlation coefficient.

### Data Analysis

Raw data files from four replicate injections each of HeLa lysate acquired at five different TIMS ramp (accumulation) times on a Bruker timsTOF Pro (3) were downloaded from ProteomeXchange (12) (PXD010012). For all searches, a protein sequence database of reviewed Human proteins (accessed 09/30/2019 from UniProt; 20463 entries including 115 common contaminant sequences) was used unless otherwise noted. Decoy sequences were generated and appended to the original database for MSFragger. PEAKS and MaxQuant only need target sequences. Tryptic cleavage specificity was applied, along with variable methionine oxidation, variable protein N-terminal acetylation, and fixed carbamidomethyl cysteine modifications. The allowed peptide length and mass ranges were 7-50 residues and 500-5000 Da, respectively. PEAKS and MaxQuant search parameters were set as close as possible to those used by MSFragger. For MSFragger searches, peptide sequence identification was performed with version 2.2 and FragPipe version 12.1 with mass calibration and parameter optimization enabled. PeptideProphet and ProteinProphet in Philosopher (version 2.0.0; https://philosopher.nesvilab.org/) were used to filter all of peptide-spectrum matches (PSMs), peptides, and proteins to 1% PSM and 1% FDR. Quantification analysis was performed with IonQuant (version 1.1.0). For PEAKS X+ searches, version 10.5 was used, and PSMs and peptides were filtered to 1% peptide FDR by clicking the FDR button on the “Summary” page. Since there is no option in PEAKS to automatically filter the proteins, we tried different protein “-10logP” scores from the smallest to the largest until the reported protein FDR was equal to 1%. MaxQuant version 1.6.10.43 was used. The PSMs and peptides were filtered to 1% PSM FDR, and the protein groups were filtered to 1% protein FDR, which are the default settings. Entries from decoy proteins and “only identified by site” were removed.

Raw data files from the mixture of three organism (5) were download from ProteomeXchange (12) (PXD014777). Three HeLa-only quality control samples (20190122_HeLa_QC_Slot1-47_1_3219.d, 20190122_HeLa_QC_Slot1-47_1_3220.d, and 20190122_HeLa_QC_Slot1-47_1_3221.d) from this same publication and repository were also used to examine gas-phase fragmentation in more recently-acquired data. In the multi-organism quantification benchmarking data set, there are two experimental conditions with three replicates each. We used MSFragger (version 2.2) coupled with FragPipe (version 12.1) and Philosopher (version 2.0.0) to perform a closed search. The protein sequence database was the combination of reviewed *H. sapiens, S. cerevisiae*, and *E. coli* proteins (accessed 04/18/2020 from UniProt; 61576 entries), with decoy sequences added. We used IonQuant (version 1.1.0) to perform quantitative analysis. For benchmarking, we downloaded MaxQuant results with the folder name “Tenzer.nomatching_MaxQuant” from https://www.ebi.ac.uk/pride/archive/projects/PXD014777. We also re-analyzed this data using MaxQuant (version 1.6.14.0) and the protein database used by MSFragger. Decoy sequences were deleted before passing it to MaxQuant. The minimum ratio count was set to 2 (default value in MaxQuant). Remaining parameters were identical to those used in the HeLa lysate analysis.

### Closed searches

Within MSFragger, precursor tolerance was set to 50 ppm and fragment tolerance was set to 20 ppm, with mass calibration and parameter optimization enabled. Two missed cleavages were allowed, and two enzymatic termini were specified. Isotope error was set to 0/1/2. 50 ppm precursor tolerance coupled with 0/1/2 isotope error encompasses deamidation (0.98 Da). Deamidated peptides are a common artifact of sample preparation and handling, so there is no need to separate these peptides from unmodified ones given the aims of this study. Additionally, this slightly wider precursor tolerance results in more candidate PSMs, which benefits expectation value estimation in MSFragger. The minimum number of fragment peaks required to include a PSM in modelling was set to two, and the minimum number required to report the match was four. The top 150 most intense peaks and a minimum of 15 fragment peaks required to search a spectrum were used as initial settings. Parameters used in PEAKS and MaxQuant were set as close as possible to those used by MSFragger.

### Semi-enzymatic searches

The parameters used by MSFragger for semi-tryptic searches were equivalent to those used in the closed searches (detailed above) but with only one enzymatic peptide terminus required. MaxQuant does not allow any missed cleavages with semi-tryptic searching. For further investigation of the identified semi-tryptic peptides, variable pyro-glutamic acid and pyro-carbamidomethyl cysteine (−17.03 Da from glutamine and cysteine), and variable water loss (−18.01) on any peptide N-terminus were also included in the semi-enzymatic MSFragger search parameters. These same parameters were used to search three HeLa injections from PXD014777 (5).

### Open searches

Precursor mass tolerance was set from -150 to +500 Da, and precursor true tolerance and fragment mass tolerance were set to 20 ppm. Mass calibration and parameter optimization were enabled. Two missed cleavages were allowed, and the number of enzymatic termini was set to two. Isotope error was set to 0. The minimum number of fragment peaks required to include a PSM in modelling was set to two, and the minimum number required to report the match was four. A minimum of 15 fragment peaks and the top 100 most intense peaks were used as initial settings.

### Label-free quantification

As there are numerous spectral preprocessing procedures, such as peak centroiding, mass calibration, and retention time alignment/calibration before peak tracing and feature extraction, tolerance settings for quantification are unlikely to translate directly between quantification tools. Thus, we decided to use the default settings for each tool, which have been optimized to perform the best in most cases. In IonQuant, mass tolerance was set to 10 ppm, retention time tolerance was set to 0.4 minutes, ion mobility (1/K_0_) tolerance was set to 0.05, normalization was enabled, and minimum isotope count was set to 2 by default. Minimum ion counts 1 and 2 were tried. In PEAKS, identification directed quantification was performed with retention time alignment, with no CV filter nor outlier removal. Mass error and ion mobility tolerances were set to 20 ppm and 0.05 1/K_0_, respectively. The retention time shift tolerance used in alignment was set to 20 minutes as recommended by the documentation. In MaxQuant, Fast LFQ was performed with large ratio stabilization, minimum ratio count set to two (except where noted), three minimum neighbors, and six average number of neighbors by default. The remaining parameters were also set to default values.

### Protein quantification with MSstats

MSstats (13) was used to calculate protein abundances from the ion abundances reported by each tool. For MSFragger and PEAKS, ions (filtered at 1% PSM and 1% protein FDR for MSFragger; 1% peptide FDR for PEAKS) were provided to MSstats. For MaxQuant, evidence.txt (filtered at 1% PSM FDR) and proteinGroup.txt (filtered at 1% protein FDR) were provided to MSstats. The dataProcess function with log10 intensity transformation was used to calculate protein abundances.

### Runtime comparisons

MSFragger (version 2.2, via FragPipe version 12.1) and MaxQuant (version 1.6.10.43) were compared on a desktop with Intel Optane SSD 900P series hard disk, Intel Core i7-8700 3.2 GHz 6 CPU cores (12 logical cores), and 32 GB memory. Due to installation and licensing constraints, PEAKS Studio X+ was used on an Intel Xeon Gold 2.4 GHz 20 CPU cores (40 logical cores) workstation with 96 GB RAM.

## Results and Discussion

### Workflow Overview

An overview of the computational workflow in shown in **Figure 1**. MS/MS spectral files acquired in PASEF mode can be read directly by MSFragger. MSFragger loads the raw format (.d) using our original spectral reading library MSFTBX (14), extended here to interact with the Bruker’s native library. During loading, Bruker’s native library (timsdata.dll or libtimsdata.so) functions are called to perform scan combining, peak picking, and de-noising. MSFTBX passes the loaded scans to MSFragger without any additional processing. After loading, MSFragger writes all extracted scans into a binary format, mzBIN, for fast data access in any future re-analyses of the same data. After database searching with MSFragger (see Experimental Procedures), PSMs are saved in the pepXML file format. PSMs are processed using PeptideProphet (15) and ProteinProphet (16) as part of the Philosopher toolkit. Philosopher is also used for FDR filtering, and for generating summary reports at the PSM, peptide ion, peptide, and protein levels (**Figure 1a**). Finally, IonQuant (see below) is used to extract peptide ion intensities for all PSMs, and adds quantification information to the PSM, peptide, and protein-level tables.

**Figure 1.**
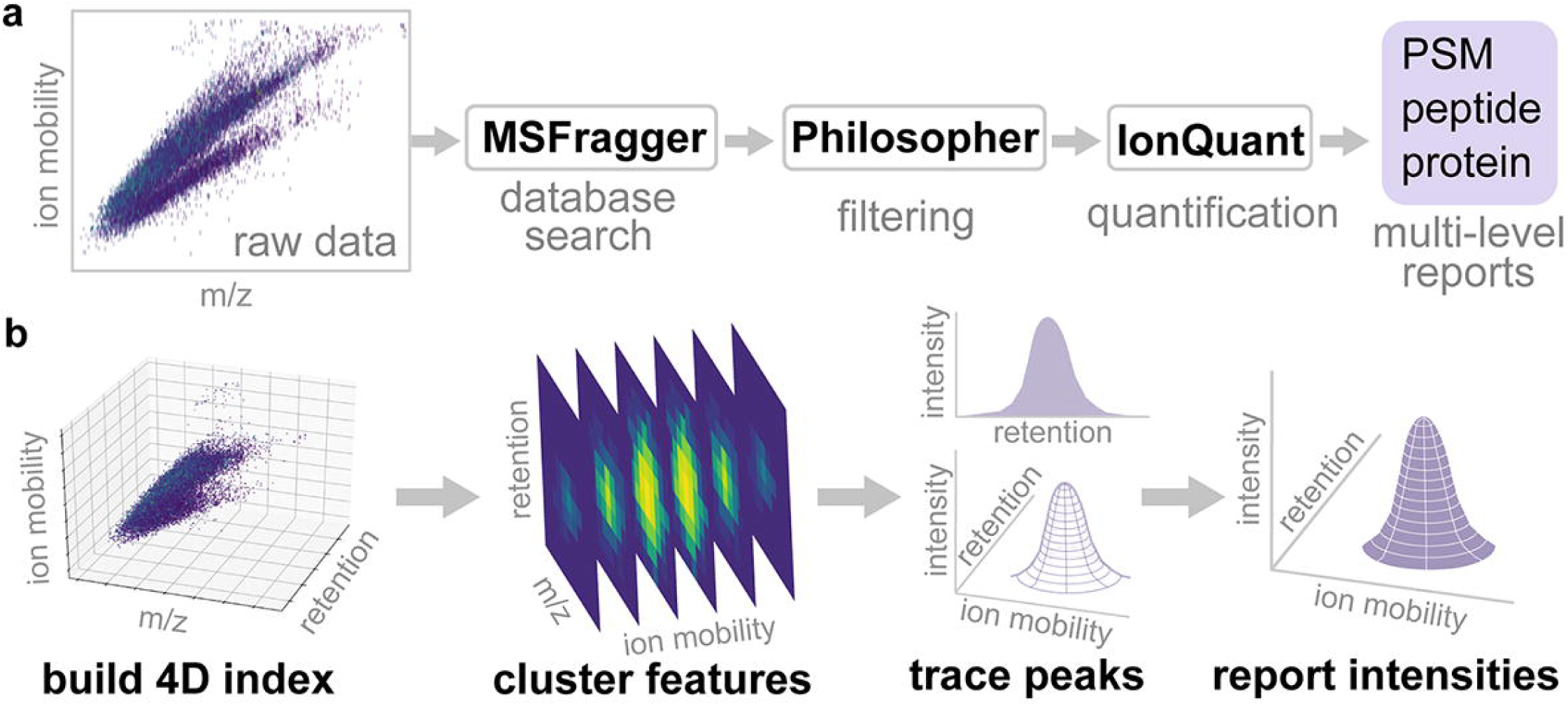
(a) Overview of the analysis workflow. Raw Bruker timsTOF data are converted (to mzBIN format) and searched with MSFragger to identify peptides from MS/MS spectra. Identifications are processed with Philosopher (PeptideProphet, ProteinProphet, FDR estimation) and FDR-filtered reports are generated at the PSM, peptide ion, peptide, and protein levels. IonQuant performs quantification and generates final reports. (b) Schematic of the IonQuant algorithm. Raw Bruker timsTOF data are loaded and indexed. Then, IonQuant traces peaks and clusters features (for all identified peptide ions) in ion mobility and retention time dimensions. Finally, IonQuant locates the apex of each peak (peptide ion) and reports its volume. Given an ion, IonQuant traces up to 3 features corresponding to 0, +1, and +2 isotopes. The total volume is reported as the ion intensity.

### IonQuant Algorithm

Spectral files generated by timsTOF PASEF are large and structurally complex due to the fast TOF scan rate and additional ion mobility dimension. IonQuant, written in Java, traces and quantifies features from the four-dimensional space (ion mobility, m/z, retention time, and intensity) quickly and accurately using indexing technology (**Figure 1b**). IonQuant first digitizes the ion mobility dimension with a predefined bin width (0.002 1/K_0_; Vs/cm^2^). Then, IonQuant indexes all peaks within this 4-D space according to their ion mobility, m/z, and retention time, which reduces memory usage and accelerates subsequent peak tracing. Given theoretical m/z, precursor ion mobility, and retention time from an identified MS/MS spectrum, IonQuant first locates the indexes corresponding to the precursor ion mobility with a user-defined tolerance. Then, it collects the m/z indexes within the tolerance of the theoretical m/z. With these two index-querying steps, IonQuant only needs to look at a small fraction of the whole data. Finally, it traverses all qualified peaks within the retention time range and generates a curve by tracing and performing Gaussian smoothing. After tracing all peaks in the retention time and m/z dimension, IonQuant traces the ion mobility dimension by clustering adjacent peaks to form 4-D features. Finally, IonQuant reports the boundaries, apex location, and volume of each detected ion feature. Given the theoretical m/z from a PSM, IonQuant tries to extract up to three 4-D features corresponding to 0, +1, and +2 isotopes. Then, it uses the summation of these features’ volumes as the quantified intensity. By default, IonQuant requires at least two isotopes (minimum isotope count 2).

IonQuant takes spectral files (.d, Bruker’s raw format, using MSFTBX as in MSFragger) and peptide identifications (pepXML) as input and outputs a csv file containing quantified results for each spectral file. When used with Philosopher summary tables as input, IonQuant adds quantification information directly to the tables containing validated PSM, peptide, and protein results. We observed that some data has nonlinear and intensity-dependent experimental errors. To get a better normalized result, we developed a piecewise normalization algorithm in IonQuant. Given all quantified runs, IonQuant first finds a “reference run” with the most ions. For each of the other runs, IonQuant calculates log-ratios of the ions overlapped with the reference run. Then, it divides the log-transformed intensities into 10 ranges. In each range, it adjusts the intensities according to the median of the log-ratios within one median absolute deviation.

In computing protein intensities from peptide ion intensities across multiple experiments, IonQuant first discards proteins with fewer quantified ions than the threshold (default 2). Then, IonQuant uses an approach similar to that of DIA-Umpire (17). Each protein’s intensity is the summed intensity of top *n* ions identified in *t* percentage of all experiments, where *n* and *t* are parameters with default values of 3 and 50%, respectively. In addition, IonQuant also uses the quantified features and the PSM table from Philosopher to generate an MSstats-compatible file for downstream analysis.

### MSFragger Has High Sensitivity in Peptide and Protein Identification

We monitored runtime and sensitivity of database searching and quantification using four replicate injections of HeLa cell digest (see **Experimental Procedures**). The data set was analyzed using MSFragger with IonQuant and compared to the results from MaxQuant and PEAKS. MSFragger identified 58954 peptides and 6525 proteins from a standard tryptic search, more than the other tools (**Table 1, Figure 2a, Supporting Table S1-S4**). Uniqueness of the peptide identifications obtained by PEAKS, MaxQuant, and MSFragger from four replicate injections of HeLa cell digest is shown in **Figure 2b**. MSFragger with IonQuant also required significantly less total analysis time than PEAKS or MaxQuant (**Figure 2c**). Furthermore, when MSFragger was used to perform subsequent searches on the same raw files (i.e. starting with mzBIN files), total processing times were under 20 minutes per file, more than nine times faster than PEAKS or MaxQuant (**Figure 2c**). We also note that a similarly fast speed can be achieved when using MGF files as input to MSFragger (generation of MGF files can be scheduled as an additional post-processing step in the instrument’s Data Analysis software immediately following data acquisition). In such a workflow, protein quantification would be limited to MS/MS-based spectral counts only, which is nevertheless sufficient for certain applications such as sample quality control or interactome analysis using affinity-purification mass spectrometry (18).

**Table 1.** Comparison of identification and quantification between the tools. Numbers of identified peptides, identified proteins, quantified proteins with/without MSstats, and median protein coefficient of variation (CV) across replicates are shown. The number of quantified proteins refers to those quantified in at least two replicates. For all searches, two missed cleavages are allowed except for MaxQuant’s semi-enzymatic search that only support zero missed cleavage. For MaxQuant, minimum 1 and 2 ratios are applied in Fast LFQ. For IonQuant, minimum 1 and 2 quantified ions are applied in native protein quantification. Since such filtering is applied in protein intensity calculation, it does not impact results from MSstats.

|  | Tool |  | Identified peptides | Identified proteins | MSstats proteins quantified | MSstats CV | Native proteins quantified | Native CV |
| --- | --- | --- | --- | --- | --- | --- | --- | --- |
| Tryptic search | PEAKS |  | 53480 | 6543 | 5227 | 0.070 | 5359 | 0.203 |
|  | MaxQuant | min 1 ratio | 44351 | 5960 | 5261 | 0.072 | 5335 | 0.072 |
|  |  | min 2 ratios |  |  |  |  | 4040 | 0.057 |
|  | MSFragger-IonQuant | min 1 ion | 58954 | 6525 | 5923 | 0.049 | 5900 | 0.073 |
|  |  | min 2 ions |  |  |  |  | 4810 | 0.064 |
| Semi-tryptic search | PEAKS |  | 87437 | 6549 | 5406 | 0.066 | 5527 | 0.194 |
|  | MaxQuant | min 1 ratio | 48606 | 5432 | 4740 | 0.072 | 4839 | 0.071 |
|  |  | min 2 ratios |  |  |  |  | 3526 | 0.054 |
|  | MSFragger-IonQuant | min 1 ion | 93466 | 6729 | 6083 | 0.046 | 6053 | 0.073 |
|  |  | min 2 ions |  |  |  |  | 4977 | 0.063 |

**Figure 2.**
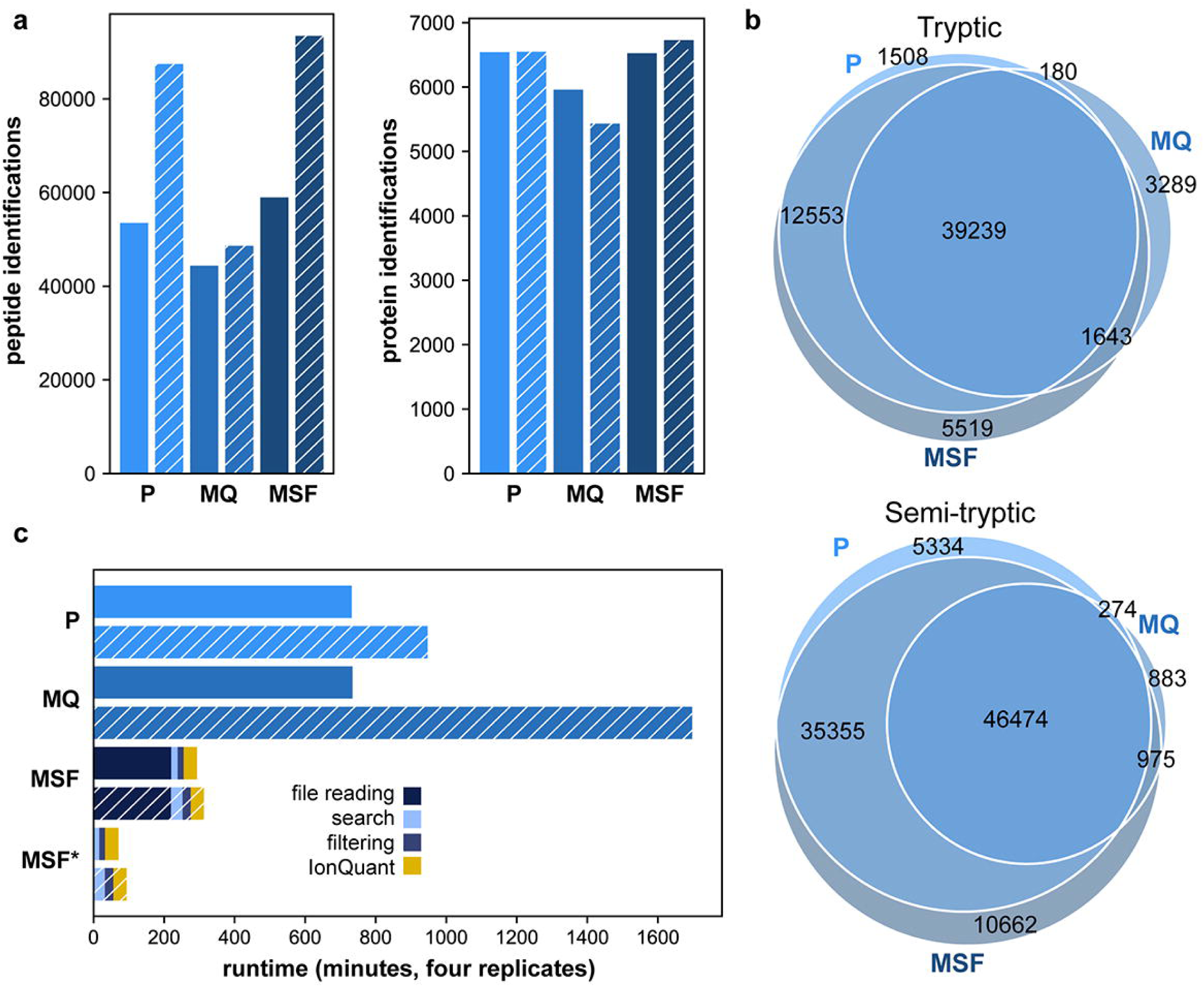
Feature identification and run time comparison. PEAKS Studio X+ (“P”), MaxQuant v1.6.10.43 (“MQ”), and MSFragger 2.2 (“MSF”) results for four HeLa replicates acquired with 100 ms accumulation time. Hatching indicates results from semi-enzymatic search. (a) Peptide (left) and protein (right) identifications. (b) Comparison of non-redundant peptide sequences identified by each tool. (c) Total analysis times for each tool. MSF* denotes MSFragger search when mzBIN files are available. MSFragger analysis times are broken down into raw file reading (i.e. conversion to mzBIN), database searching, filtering, and quantification with IonQuant.

### Precise Protein Quantification with IonQuant

We evaluated the quantitative performance of MSFragger with IonQuant and compared with MaxQuant and PEAKS, using the tryptic search results (see **Experimental Procedures**) from the same four HeLa replicates (**Table 1**). Because each tool groups peptides and performs protein quantification differently, we used MSstats to independently calculate protein abundances from ions quantified by these tools. Across the four replicate injections, IonQuant with MSstats demonstrated excellent reproducibility, with Pearson correlation between replicates of 0.97 or above (**Figure 3a**), higher than that from PEAKS and MaxQuant (**Supporting Figure S1**). The distribution of CVs for each protein among the tools is shown in **Figure 3b**. Considering proteins quantified in at least two replicates, IonQuant with MSstats quantified the most proteins (5923) while exhibiting the smallest median CV across replicates of 0.049, compared to PEAKS-MSstats (0.070) and MaxQuant-MSstats (0.072). Protein abundances reported by IonQuant correlated with those reported by PEAKS and MaxQuant with Pearson correlations of 0.84 and 0.83, respectively (**Figure 3c, Supporting Figure S2 shows the ion-level correlation**). Each tool, including IonQuant, can also perform peptide to protein roll-up and report protein-level quantification (‘native’ quantification in **Table 1**). However, our analysis shows that post-processing using MSstats performed as well as or better than native protein-level quantification methods for all three tools. For MaxQuant, applying an additional filter of 2 minimum peptides per protein for quantification (the default setting in MaxQuant) reduced the mean protein CV to 0.057. However, this was associated with a very significant drop in the total number of proteins quantified in at least two replicates (from 5335 to 4040, **Table 1**). IonQuant has a similar option (minimum ion count parameter) for its native quantification method. Requiring at least 2 quantified ions per protein, the median protein CV reduced to 0.064 with a corresponding reduction in the number of quantified proteins to 4810 (**Table 1**).

**Figure 3.**
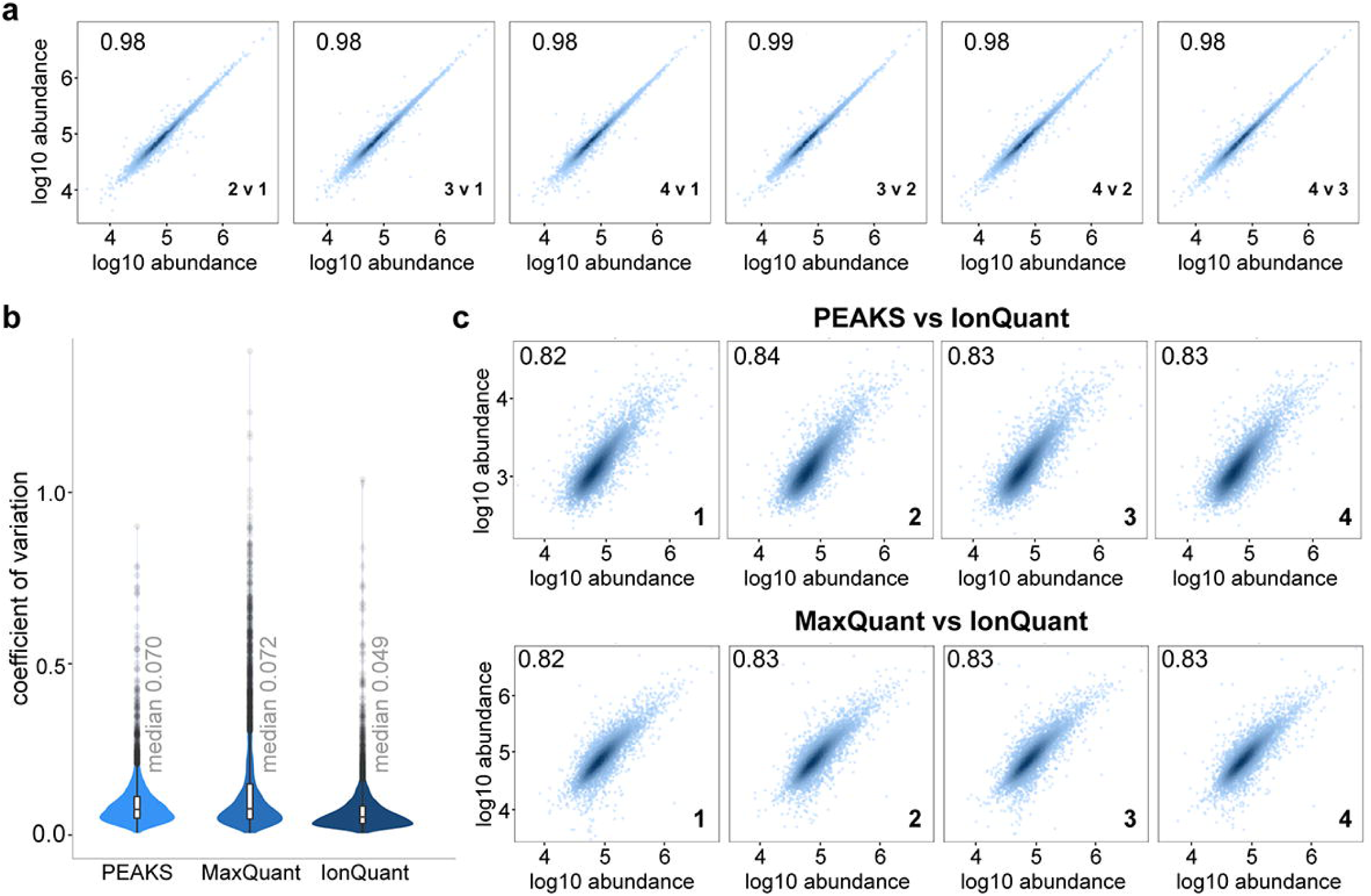
Protein quantification (with MSstats). (a) Correlation of quantified proteins between four technical replicates, MSFragger-IonQuant results. Each paired comparison is labeled in the bottom right-hand corner of the plot. (b) Protein coefficient of variation across the four replicates, comparing PEAKS, MaxQuant, and MSFragger-IonQuant. Replicates are labeled in the bottom right-hand corner of each plot. (c) Comparison of MSFragger-IonQuant protein abundances to PEAKS and MaxQuant for each replicate. Log-transformed intensities from IonQuant are shown on the x-axis.

### Quantification Accuracy Evaluation Using Three Organism Data

After showing the good precision (CV across replicates) of IonQuant, we used data from the mixture of three organisms (*H. sapiens, S. cerevisiae*, and *E. coli*) published by Prianichnikov et al. (5) to evaluate our tool’s accuracy. The data set consists of two samples (A and B) with different amounts of each proteome, such that the resulting ratios between A and B are 1:1 (*H. sapiens*), 2:1 (*S. cerevisiae*), and 1:4 (*E. coli*). There are three technical replicates of each sample. We first performed a closed search on these data using MSFragger (version 2.2) coupled with FragPipe (version 12.1) and Philosopher (version 2.0.0), then quantified using IonQuant (version 1.1.0), trying both minimum ions set to 1 and 2. We also used MSstats to calculate the protein intensity as a comparison. MaxQuant results as provided by the authors are used as the benchmark. After removing decoy proteins and those “only identified by site” there are 4369 proteins quantified by MaxQuant in both experimental conditions. We also reanalyzed the data using MaxQuant (version 1.6.14.0) with the protein database used by MSFragger and minimum ratio count 2, and obtained similar results (a total of 4454 proteins were quantified in both conditions).

We used LFQBench (19) to evaluate the data and plot the figures without applying any additional filtering. In **Figure 4**, *S. cerevisiae* proteins are shown in orange, *H. sapiens* in green, and *E. coli* in purple. Box plots to the right of each scatter plot show the distribution of the protein intensities. Both IonQuant and MaxQuant could recover the ratios between organisms well, but IonQuant quantified more proteins with the same minimum ion/ratio count of 2. With a minimum ion count of 1, IonQuant quantified significantly more proteins (6582 compared to 4890 with minimum ion count 2), albeit with an increased number of outliers. Since the minimum ion count filtering is only applied to native protein intensity calculation, the number of proteins from MSstats is close to native protein quantification with minimum ion count 1. MSstats, however, results in fewer outliers than native IonQuant method due to its more advanced peptide to protein roll-up algorithm.

**Figure 4.**
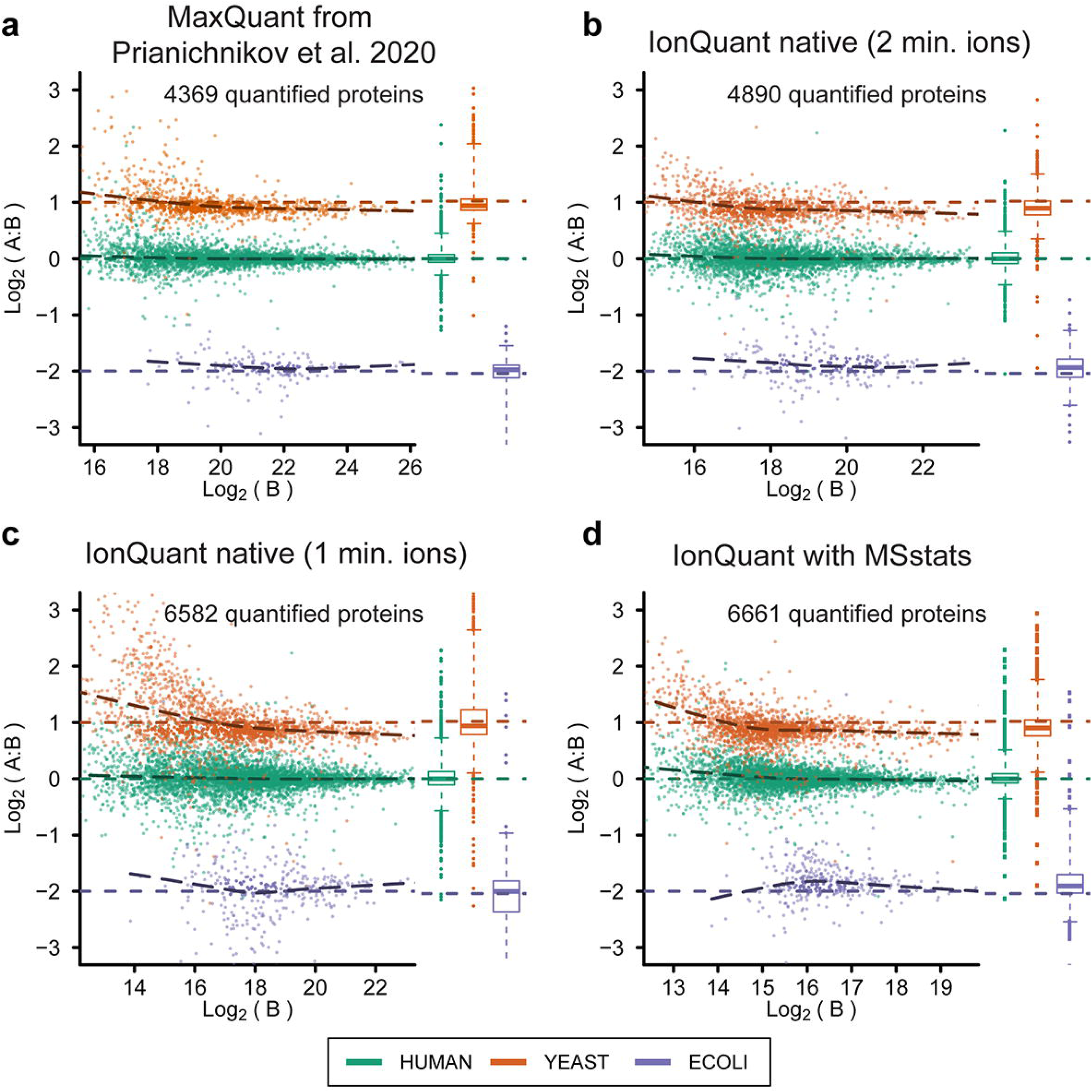
Protein intensities from IonQuant and MaxQuant from the three organism benchmarking data set. *S. cerevisiae* proteins are shown in orange, *H. sapiens* in green, *E. coli* in purple. The ground truth ratios are shown in the colored dashed lines (2:1, 1:1, and 1:4, for *S. cerevisiae, H. sapiens*, and *E. coli* respectively). Box plots on the right of each scatter plot show the distributions of the intensities for each organism. (a) MaxQuant result published by Prianichnikov et al. 2020. 2. (b) IonQuant result with minimum ion count equals 2. (c) IonQuant result with minimum ion count equals 1. (d) IonQuant result with MSstats calculating the protein intensity.

### Open Search Analysis

Using MSFragger and IonQuant, we performed a quantitative open search on the four HeLa replicates acquired with 100 ms accumulation time. After statistical evaluation and filtering by Philosopher, mass shifts corresponding to water and ammonia losses (−17 and -18 Da, respectively) were the most prominent, followed by a +52.91 Da mass shift that corresponds to substitution of three protons with Fe(III), possibly an artifact from sample handling. Open search also revealed the presence of many semi-tryptic (neutral loss) peptides. Plots displaying the number of PSMs for each of these mass shifts are shown in **Supporting Figure S3 (Supporting Table S5-S6)**. MSFragger and IonQuant analysis times were not significantly longer for open search.

### Semi-tryptic Peptide Monitoring

From the open search, we observed a significant number of semi-tryptic PSMs, and PSMs with water and ammonia loss. Intrigued by these observations, we investigated whether this was indicative of ion activation prior to MS/MS analysis. To this end, we performed semi-enzymatic searches (also allowing -17 and -18 Da losses, see **Experimental Procedures**) on the HeLa data acquired with different TIMS accumulation times (3), during which trapping in the first TIMS region and mobility separation in the second occur. Across the five different accumulation times tested in the publication (25, 50, 100, 150, and 200 ms), we observed that the number of PSMs with only one enzymatic terminus increases with accumulation time (**Figure 5a**). The relationship between accumulation time and semi-tryptic peptides is likely due in part to increased sensitivity. The number of peptide ions that can be targeted for fragmentation increases with accumulation time (3), so low-intensity ions are more likely to be detected when longer accumulation times are used. This can be seen in **Figure 5b**, where the share of total ion intensity from semi-tryptic peptides increases as the instrument has more time to interrogate these lower-abundance ions. We also monitored the abundance ratio of each tryptic peptide to its corresponding semi-tryptic peptide and found the same trend across accumulation time reflected in this pairwise comparison (**Supporting Figure S4**).

**Figure 5.**
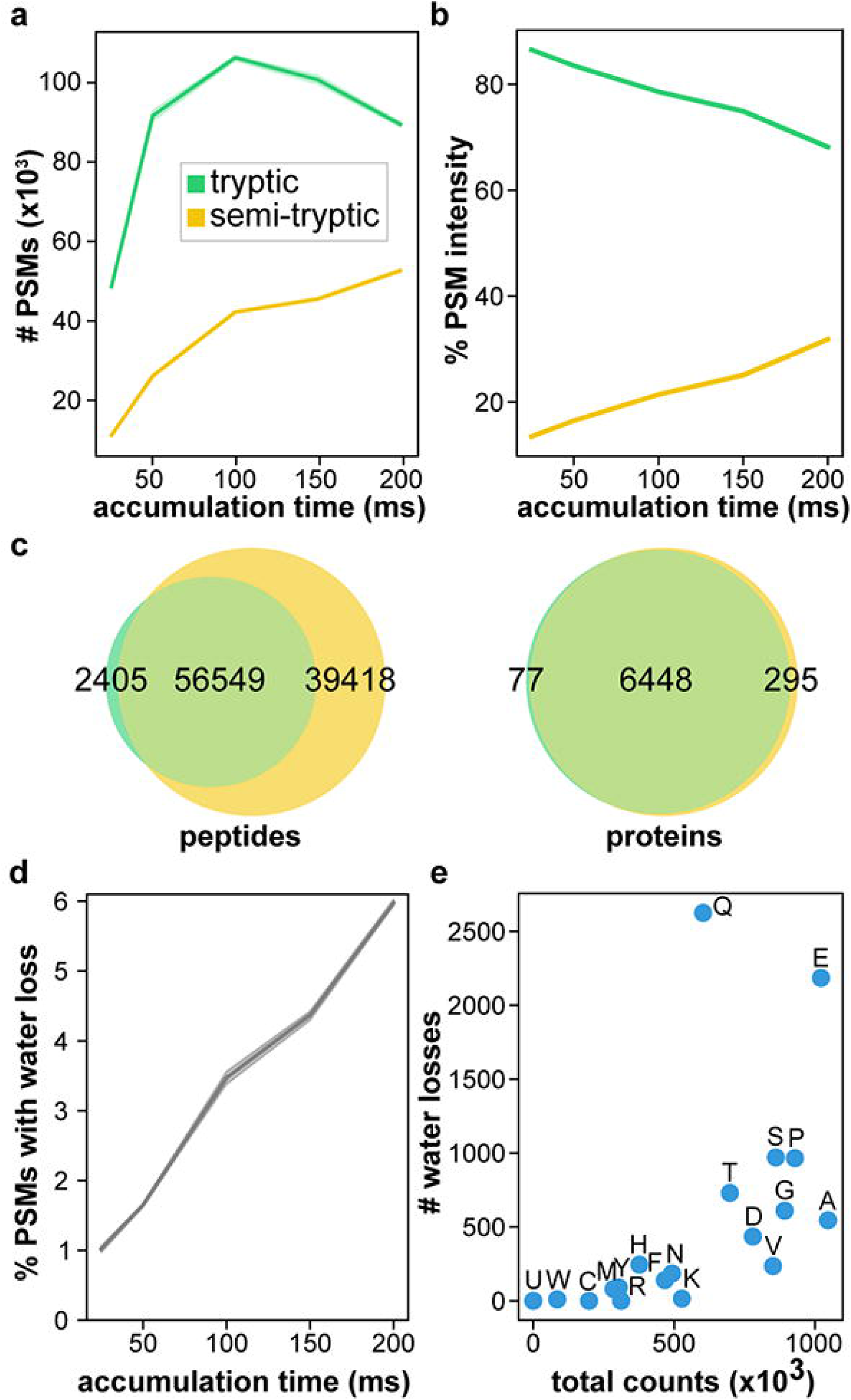
Semi-tryptic searching with MSFragger monitors fragmentation within dual TIMS device. The total number of semi-tryptic PSMs (a) and the percentage of total precursor intensity from semi-tryptic PSMs (b) increase with accumulation time. (c) More peptides and proteins are identified using semi-tryptic search with MSFragger (four pooled HeLa replicates, 100 ms accumulation time). For semi-tryptic search, variable pyro-glutamic acid and pyro-carbamidomethyl cysteine (−17.03 Da from glutamine and cysteine), and variable water loss (−18.01) allowed on any peptide N-terminus were added. (d) The percentage of PSMs displaying neutral water loss increases with accumulation time. (e) Water losses for each amino acid following the cleavage site are plotted against the total occurrences of the amino acid in the data set. For each line plot, shaded areas represent the 95% confidence interval from four replicates.

At 100 ms accumulation time, which was selected as optimal by the authors of the original manuscript, a semi-enzymatic MSFragger search resulted in an astonishing ∼60% increase in the number of identified peptides (from 58954 to 93466 across four replicates, or 95967 with ammonia and water losses as variable modifications), counting overlapping semi-tryptic and fully tryptic peptides as different peptides (**Figure 2** and **Figure 5c**). The number of identified proteins from the MSFragger search increased as well (from 6525 to 6729, or 6749 with ammonia and water losses as variable modifications). Both PEAKS and MSFragger identified more unique peptides with a semi-enzymatic search (**Figure 2a, b**), PEAKS identified ∼63% more and MSFragger identified ∼59% more, while MaxQuant results did not reflect a noticeable increase. This may be partially attributed to the fact that MaxQuant does not allow missed cleavages in semi-enzymatic searches. Of those peptides with a single enzymatic terminus identified by the semi-enzymatic search, the majority (67%) were found alongside their full-length tryptic form. We also demonstrate that MSFragger with IonQuant quantifies more proteins in semi-tryptic vs. tryptic search without compromising accuracy (**Table 1**). It is also worth noting that, due to fast fragment ion indexing (less than 10 seconds for closed and open tryptic searches, maximum of 80 seconds for semi-tryptic searches), MSFragger’s runtime advantage over MaxQuant and PEAKS is even greater when performing more complex search tasks, such as semi-enzymatic searches (**Figure 2c**).

We further investigated the source of observed semi-tryptic peptides. We compared the apex retention times of unmodified semi-tryptic peptides to their corresponding fully-tryptic peptide and found that, across the entire data set (four replicates each of five different accumulation times), 76% were within 60 seconds of one another, indicating that these semi-tryptic peptides largely originated within the instrument. Among all identified semi-tryptic peptides, proline was most likely to be found C-terminal to the cleavage site, consistent with known fragmentation behavior of positively-charged peptides (20, 21). Furthermore, in the semi-enzymatic searches, we allowed a neutral loss of H_2_O from any N-terminal residue. We observed an increase in the percentage of PSMs containing a water loss with longer accumulation times (**Figure 5d**), as would be expected for a gas-phase fragmentation event. As described previously (22-25), water loss from N-terminal glutamine and glutamate is frequently observed following collision-induced dissociation (CID) of peptides. Of the peptides identified with N-terminal semi-tryptic cleavages, we observed that water loss occurred preferentially when glutamine or glutamate were present C-terminal to the cleavage site (**Figure 5e**). As the semi-tryptic peptides identified in this data set display neutral losses characteristic of CID, it appears peptide ion activation occurred in the dual TIMS device, resulting in the majority of the semi-tryptic peptides we observe.

The high rates of semi-tryptic PSMs may be specific to the timsTOF data sets used in this work, and these analyses should be repeated as more data sets become publicly available. In general, we expect improvements in instrument tuning to provide gentler peptide ion handling and therefore less fragmentation within the instrument. Indeed, when examining three replicate injections of HeLa digest (60100 ± 200 unique peptide identifications on average) from a recently published timsTOF PASEF DDA data set (5), we find that the percentage of semi-tryptic peptides decreases, from 28% to 17%, when the same accumulation time (100 ms) is used. For certain applications, e.g. in HLA peptidome profiling studies that require precise characterization of peptide sequences (26, 27), further reduction in in-TIMS fragmentation with altered tuning settings may be necessary. On the other hand, reducing the energy imparted by the source and initial ion optics can reduce ion transmission, in some cases dramatically. In many analyses it may thus be preferable to use higher energies in the instrument source (or later ion optics such as the TIMS device itself) to improve transmission efficiency despite increased fragmentation of some peptides, making a semi-enzymatic search necessary to recover the identities of all peptides analyzed (28) and maximize the sensitivity of the instrument. Furthermore, certain analyses, such as those of glycopeptides (29) may actually benefit from in-source pseudo-MS^3^ capabilities to enable advanced acquisition methods. As the in-TIMS fragmentation level seems to be tunable, the instrument appears to have the capability to perform these pseudo-MS^3^ methods as well.

### Spectral Library Generation

The search results from MSFragger (after processing with Philosopher/PeptideProphet) can also be fed into Skyline (30) to generate spectral libraries and inspect peptide features in three dimensions (**Supporting Figure S5**). Skyline can also be used to perform MS1-based quantification, as well as targeted quantification from data independent acquisition (DIA, diaPASEF) data (31). By providing Skyline with 1% FDR filtered protein list (generated by Philosopher, in FASTA format), Skyline libraries can be effectively created with desired protein level and peptide ion FDR filters (e.g. 1% protein FDR and 1% peptide ion FDR). A detailed tutorial for importing and visualizing the results from MSFragger search in Skyline can be found on the MSFragger webpage (https://msfragger.nesvilab.org/tutorial_pasef_skyline.html). Furthermore, the spectral library building tool EasyPQP (https://github.com/grosenberger/easypqp) has been adapted to be used with ion mobility data, and we incorporated this capability into the MSFragger user interface FragPipe. This feature allows building spectral libraries from DDA data as part of DIA workflows, e.g. for subsequent quantification from DIA data using OpenSWATH (32), Spectronaut (19), or DIA-NN (33) (limited support for diaPASEF data at the time of writing). Running EasyPQP on MSFragger tryptic search results of the four HeLa replicates (100 ms accumulation time) resulted in a spectral library containing 58931 peptides.

### Conclusions

Due to the efficient parallel accumulation strategy and the added selectivity of trapped ion mobility, the timsTOF PASEF method has achieved highly sensitive proteomics measurements. We have extended MSFragger to directly read raw PASEF data for rapid database searching, and developed IonQuant to accurately quantify peptides and proteins from these data. For standard tryptic searches, MSFragger requires less than half the analysis time needed by other tools that currently support PASEF data, and is three to five times faster for semi-enzymatic searching while still annotating the greatest number of peptides among the tools compared. MSFragger is the only PASEF-compatible search engine with the ability to conduct open searches in reasonable time. The flexibility afforded by MSFragger’s modest analysis times can be applied for post-translational modification (PTM) discovery or screening for artifacts of sample preparation or data acquisition. Overall, we report data analysis times two- to five-fold shorter than existing tools that remove a primary bottleneck in the usability of timsTOF PASEF data. MSFragger and IonQuant enable fast, sensitive, and precise quantitative proteomic analyses, including semi-enzymatic and open searches, as well as spectral library generation for diaPASEF analysis workflows. A match-between-runs (MBR) capability for IonQuant, including MBR FDR control, is under development and will be described in future work. This entire pipeline can be accessed through a graphical user interface FragPipe (http://fragpipe.nesvilab.org/) or with the command line for high-throughput applications. Outputs are also compatible with tools such as Skyline, MSstats, and with proteomics data viewer PDV (34) for visualization of peptide assignments to MS/MS spectra, enabling a variety of complete workflows.

## Abbreviations

LC-MS: liquid chromatography-mass spectrometry
TIMS: trapped ion mobility spectrometry
TOF: time-of-flight
PASEF: parallel accumulation-serial fragmentation
DDA: data-dependent acquisition
DIA: data-independent acquisition
MS/MS: tandem mass spectrometry
PSM: peptide-spectrum match
LC-IMS-MS: liquid chromatography-ion mobility-mass spectrometry
XIC: extracted ion chromatogram
CV: coefficient of variation
LFQ: label free quantification
FDR: False Discovery Rate
CID: collision-induced dissociation
CPU: central processing unit
PTM: post-translational modification
MBR: match-between-runs

## Acknowledgements

The authors would like to thank Markus Lubeck and Florian Meier for helpful discussions, and George Rosenberger for assistance with EasyPQP. We also thank the users of our tools for their feedback. This work was funded in part by NIH grants R01-GM-094231 and U24-CA210967.

## Data and Software Availability

The HeLa-only data used in the manuscript were published by Meier et al. (3) and can be found from the ProteomeXchange Consortium via the PRIDE partner repository (35) with the identifier PXD010012 (https://www.ebi.ac.uk/pride/archive/). The *S. cerevisiae - H. sapiens - E. coli* data were published by Prianichnikov et al. (5) and can be found under the ProteomeXchange identifier PXD014777. MSFragger and IonQuant programs were developed in the cross-platform Java language and can be accessed at http://msfragger.nesvilab.org/ and https://github.com/Nesvilab/IonQuant. The DOI of the IonQuant tool used in this manuscript (version 1.1.0) is 10.5281/zenodo.3828088.

## Author Contributions

F.Y. adopted MSFragger for timsTOF PASEF data and developed the IonQuant algorithm, with contribution from G.C.T. and D.A.; S.E.H., F.Y., and A.I.N. analyzed the data; S.E.H., F.Y., D.A.P., and A.I.N. wrote the manuscript with input from all authors; A.I.N. supervised the entire project.

