## Supplemental Figure for "Fast quantitative analysis of timsTOF PASEF data with MSFragger and IonQuant"

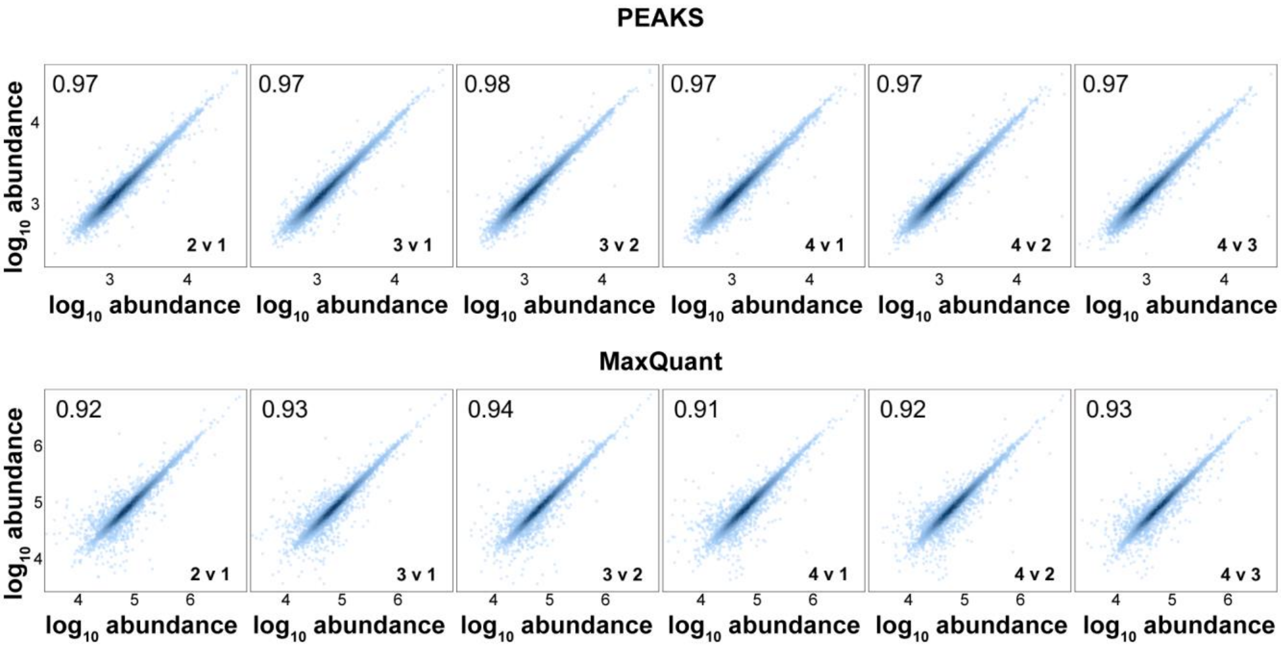


**Figure S1.** PEAKS and MaxQuant protein quantification compared between replicates. Comparisons are labeled in the lower right corner of each plot (e.g. '2 v 1', where protein abundances from replicate 1 are on the x-axis, and those from replicate 2 are plotted on the y-axis). Pearson correlation coefficients are shown in the upper left corner.

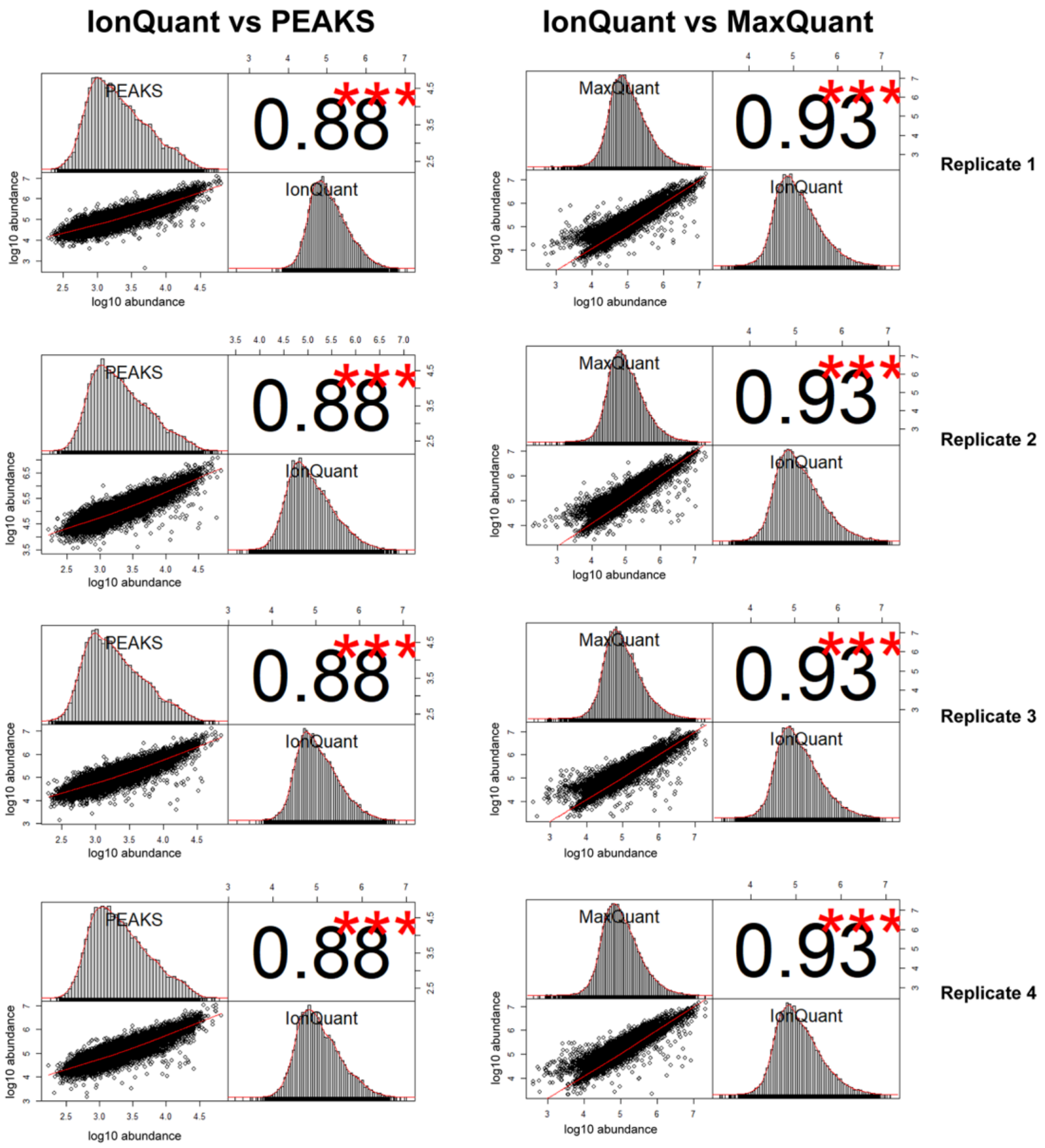


**Figure S2.** PEAKS (left column) and MaxQuant (right column) ion quantification compared to IonQuant for each replicate. Within each scatterplot, IonQuant ion abundances are plotted on the y-axis. Pearson correlation coefficients are shown in the upper right quadrant of each plot, and histograms of peptide ion abundances are shown for each tool. There are two bands in the scatter plot with MaxQuant. After careful examination, we attribute this to differences in treatment of multiply-charged ions. For singly-charged ions, IonQuant and MaxQuant have a good correlation (0.98 Pearson correlation, data not shown).

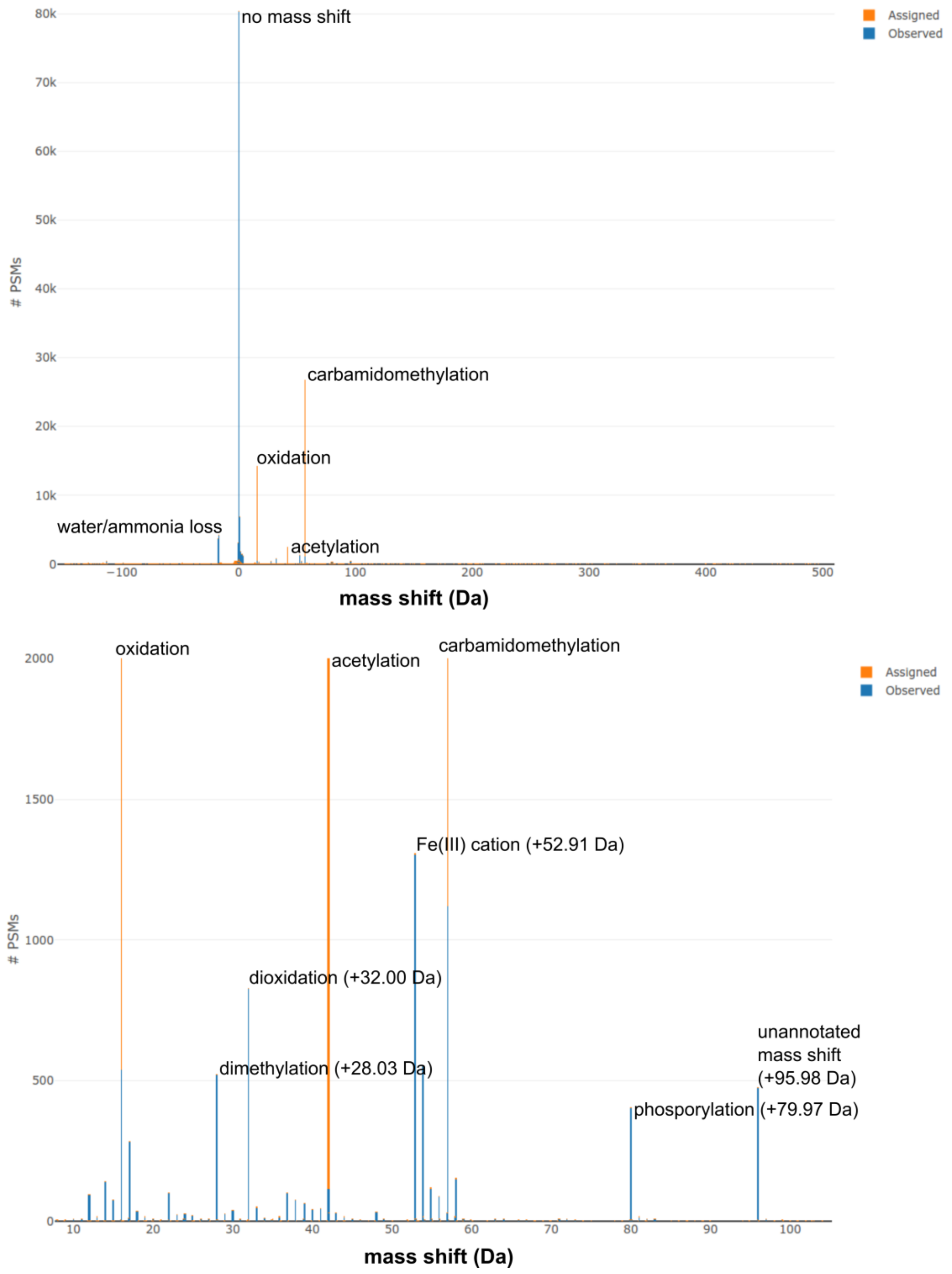


**Figure S3.** Mass shifts found by open searching one HeLa injection acquired using 100 ms accumulation time with MSFragger. Top, PSM count is shown for the full mass range (-150 to +500 Da). Bottom, detail of +10 to +100 Da (y-axis is also zoomed). Peaks in blue ('Observed') are annotated with likely modifications from Unimod, orange peaks ('Assigned') were those specified as variable modifications in the search.

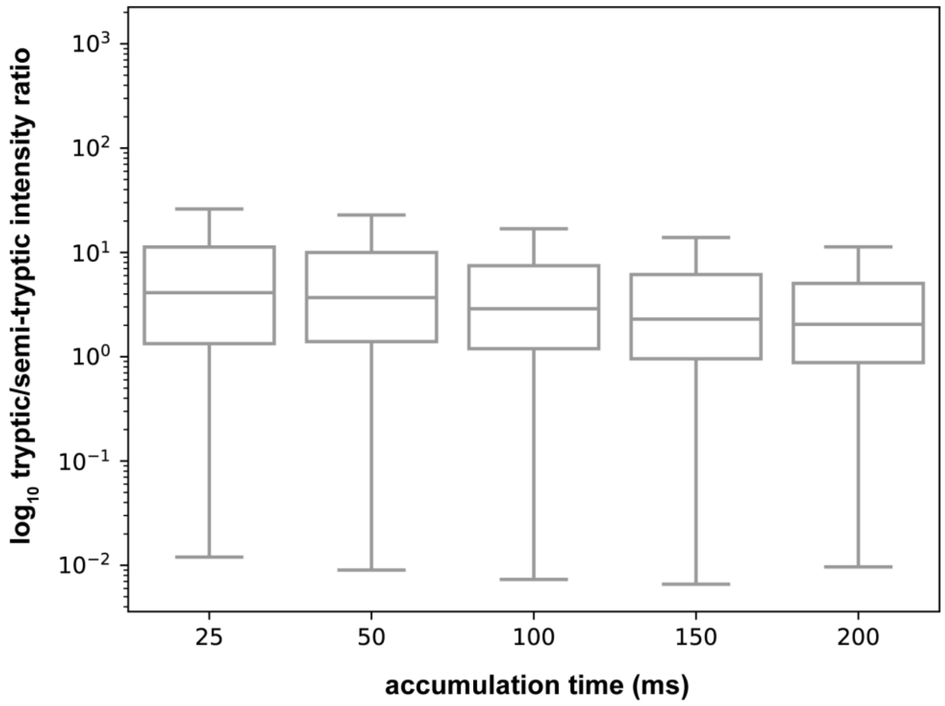


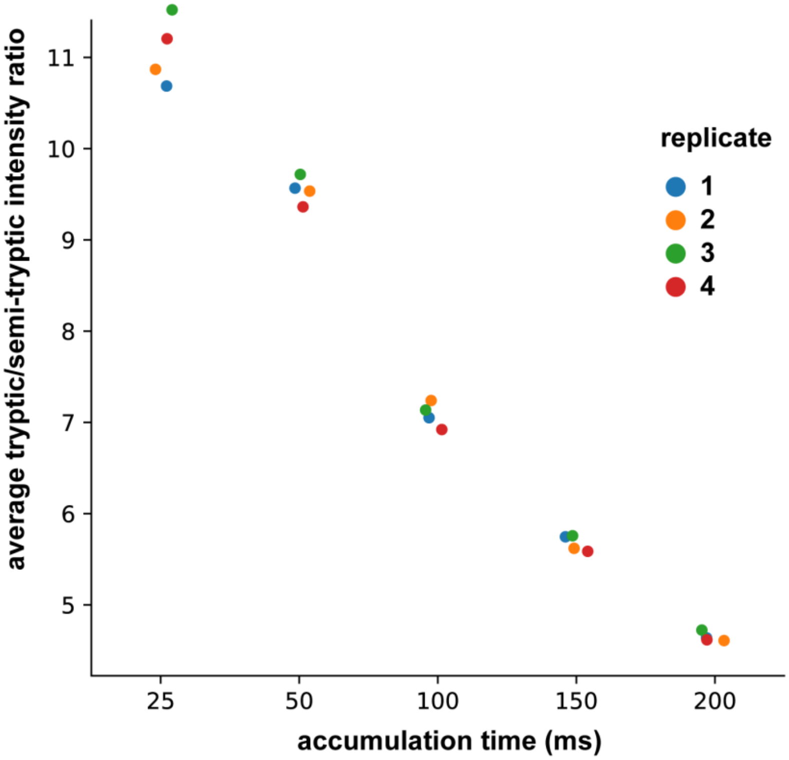


**Figure S4.** Longer accumulation time results in lower ratio of intact peptide ion intensity to its corresponding semi-tryptic fragment. Ratios of unmodified tryptic to semi-tryptic apex intensities are shown across the five accumulation times (top). Average tryptic/semi-tryptic abundance ratios for each replicate are shown across the five accumulation times (bottom).

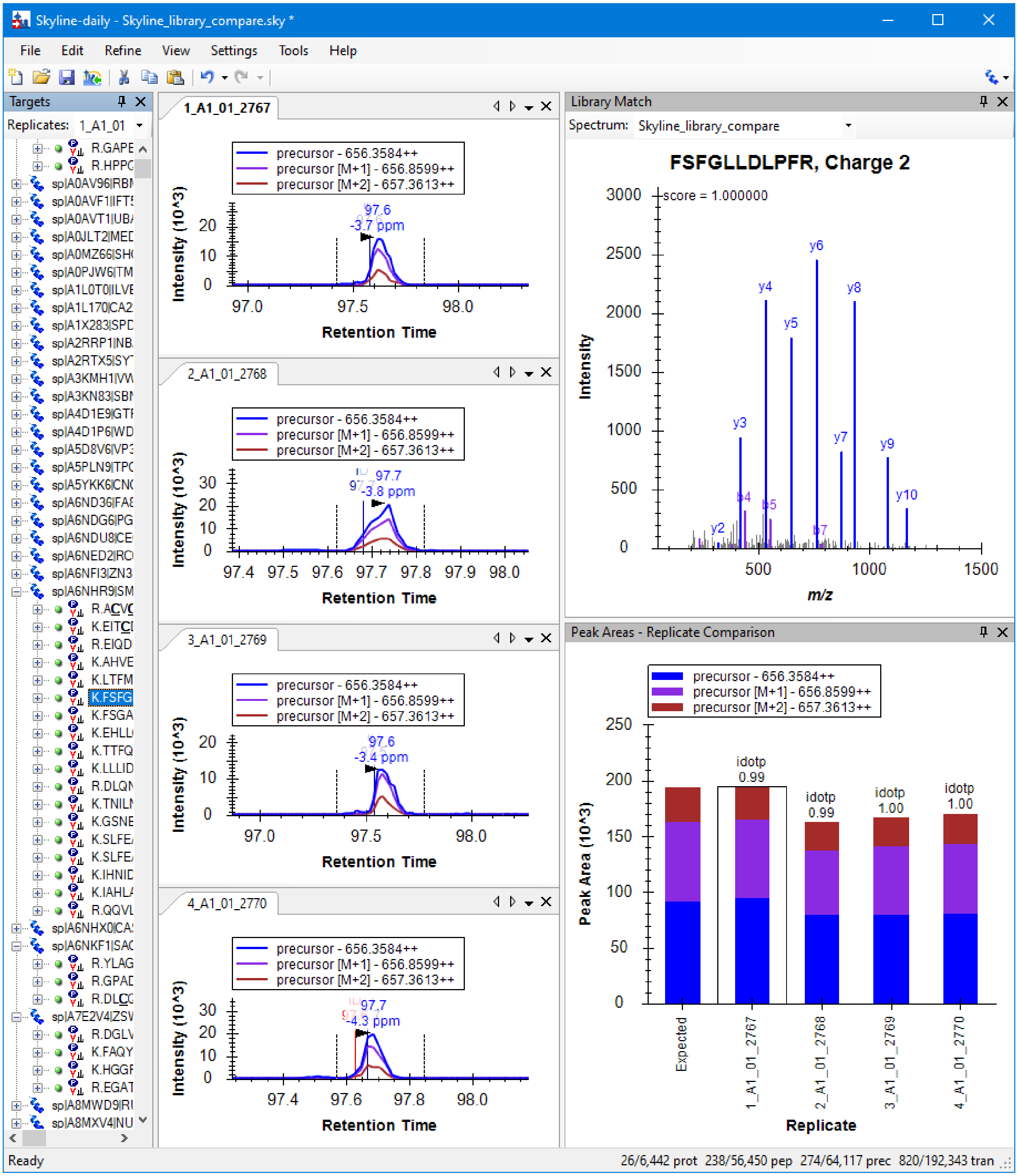


**Figure S5.** MSFragger results viewed in Skyline. Chromatograms and peak areas for the same peptide in four different replicates are shown after feature extraction using the Skyline spectral library.
